# AMPK regulates cell shape of cardiomyocytes by modulating turnover of microtubules through CLIP-170

**DOI:** 10.1101/2020.05.29.123299

**Authors:** Shohei Yashirogi, Toru Katayama, Takemasa Nagao, Yuya Nishida, Hidetaka Kioka, Tsubasa S Matsui, Shigeyoshi Saito, Yuki Masumura, Osamu Tsukamoto, Hisakazu Kato, Issei Yazawa, Hiromichi Ueda, Osamu Yamaguchi, Kenta Yashiro, Satoru Yamazaki, Seiji Takashima, Yasunori Shintani

## Abstract

AMP-activated protein kinase (AMPK) is a multifunctional kinase that regulates microtubule (MT) dynamic instability through CLIP-170 phosphorylation; however, its physiological relevance in vivo remains to be elucidated. In this study, we identified an active form of AMPK localized at the intercalated discs in the heart, a specific cell-cell junction present between cardiomyocytes. A contractile inhibitor, MYK-461, prevented the localization of AMPK at the intercalated discs, and the effect was reversed by the removal of MYK-461, suggesting that the localization of AMPK is regulated by mechanical stress. Time-lapse imaging analysis revealed that the inhibition of CLIP-170 Ser-311 phosphorylation by AMPK leads to the accumulation of MTs at the intercalated discs. Interestingly, MYK-461 increased the individual cell area of cardiomyocytes in CLIP-170 phosphorylation-dependent manner. Moreover, heart-specific CLIP-170 S311A transgenic mice demonstrated elongation of cardiomyocytes along with accumulated MTs, leading to progressive decline in cardiac contraction. In conclusion, these findings suggest that AMPK regulates the cell shape and aspect ratio of cardiomyocytes by modulating the turnover of MTs through homeostatic phosphorylation of CLIP-170 at the intercalated discs.

## Introduction

AMP-activated protein kinase (AMPK) can sense the increase of intracellular AMP or ADP concentration, and is fully activated by the phosphorylation of conserved Thr residue in the activation loop by upstream kinases, including LKB1 or Ca^2+^/calmodulin-activated protein kinase kinases, CaMKK2 [1]. Canonical stimulation known to activate AMPK is energetic stress, and this explains why AMPK switches on downstream signaling pathways involved in ATP production while switching off the anabolic pathways. However, AMPK can be activated by various stimuli other than energetic stresses including Ca^2+^ increase, oxidative stress, or genotoxic stress. The downstream effects of AMPK are not just restricted to the regulation of metabolism. It has also been demonstrated that AMPK is a multifunctional kinase known to regulate cell cycle, polarity, membrane excitability, and a variety of cellular functions by phosphorylating specific sets of substrates, presumably in a spatiotemporal manner [1–3].

As such, we previously demonstrated that AMPK controls directional cell migration by modulating microtubule (MT) dynamic instability through direct phosphorylation at CLIP-170 Ser-311 in Vero cells [4]. MTs are noncovalent polymers comprised of tubulin heterodimers, and one of the major constituents of cytoskeleton. Although the term of cytoskeleton suggests static structure, MTs are in fact highly dynamic, especially the plus end of MTs, which exhibits a behavior called as dynamic instability; individual MT ends fluctuate between polymerization and depolymerization phase [5]. CLIP-170 is one of the MT plus end tracking proteins (+TIPs), and its role is to bind MT plus end in order to protect them from depolymerizing factors, thereby MT polymerization is accelerated. Conversely, MT depolymerization is promoted when CLIP-170 departs from plus end [6,7]. This dissociation of CLIP-170 has been shown to be regulated by AMPK-mediated phosphorylation of CLIP-170 Ser 311 residue [8,9]. In migrating Vero cells, the inhibition of AMPK or the expression of CLIP-170 Ser 311-to-Ala mutant (CLIP-170 S311A), which is a non-phosphorylatable mutant, leads to an increase in the amount of stable MTs and disturbed cell polarity, thereby resulting in the impairment of free or directional cell migration [4]. We originally found CLIP-170 as a novel substrate of AMPK from mouse heart homogenates, the relevance of AMPK-CLIP-170 on MT dynamic instability in vivo, however, remains to be elucidated.

One of the most important findings regarding cardiac MTs is that density of MTs increases during end-stage heart failure regardless of their etiology [10,11]. In a mouse pressure-overload heart failure model, the expression of MTs in the heart was shown to be increased [12]. Interestingly, the treatment with MT depolymerizer, colchicine, reversed the MT accumulation and improved cardiac function and survival rate [13]. Conversely, anti-cancer drug, paclitaxel, which has been shown to stabilize MTs, was reported to induce cardiac dysfunction as a side effect [14]. Moreover, increasing MT stability impairs contraction and thus is associated with human heart failure [15]. These findings suggest that effective reversal of cardiac MT stability will have therapeutic potential for the treatment of heart failure [11,16]. However, upstream and downstream mechanisms of MT stabilization in vivo heart are still not well understood.

As an integral component of cytoskeleton, cardiac MTs are also important for the maintenance of cell shape, that is, aspect ratio (length/width ratio). Cardiomyocytes adapt to elasticity of the extracellular matrix and modulate their aspect ratio in such manner that they can maximize its systolic performance. When cardiomyocytes changes its aspect ratio on stiff gels, MT polymerization increases, however, other cytoskeleton components are not involved in changing the cell shape [17].

Cardiomyocytes exhibit unique property of cellular polarity through which they are connected to the neighboring cardiomyocytes only at the short side of the cells, which is referred to as the intercalated disc. The intercalated disc is a special form of cell-cell junctions in cardiomyocytes, which consists of three types of cell-cell junctions, namely adherence junctions, gap junctions, and desmosomes. They cooperatively reinforce the synchronized cardiac contraction by producing of mechanical stability, transmission of forces generated by myofibrils, and electrical coupling [18,19]. Furthermore, it has been demonstrated that a transient receptor potential, vanilloid family type 2 (TRPV2) cation channel localizes at the intercalated discs and serves as a mechanoreceptor to maintain cardiac structure and function [20]. The structure of intercalated discs modulate in response to hemodynamic stress; therefore, mechano-signaling has to be involved in such feedback system [18,19]. Although the intercalated discs play an important role in cardiac homeostasis and pathophysiology, the mechanism of their maintenance in cardiomyocytes and details of mechano-signaling initiated from them are largely unknown.

In this study, we revealed that AMPK is localized at the intercalated disc in the heart, where it regulates MT dynamics through CLIP-170 phosphorylation. The localization of AMPK was regulated by mechanical stress. Inhibition of CLIP-170 phosphorylation resulted in the accumulation of MTs and an increase in individual cell area. Our data also revealed the important association between mechano-signaling and regulation of cell shape through MT dynamics, which is regulated by AMPK at the intercalated discs.

## Results

### AMPK and phosphorylated CLIP-170 are localized at the intercalated discs in murine heart

In order to gain an insight into the relevance of AMPK and CLIP-170 in the heart, we first assessed and compared phosphorylation levels of AMPK and CLIP-170 in both heart and liver in different developmental stages from embryonic day 15 to 8 weeks after the birth of mouse (Fig. 1A and B). In the liver, the phosphorylation levels of AMPK and acetyl-CoA carboxylase (ACC), which is a crucial substrate of AMPK known to be involved in the regulation of metabolism, increased simultaneously with mouse development, implying an important role of AMPK in the regulation of systemic metabolism. Conversely, phosphorylation levels of CLIP-170 were not changed in the liver throughout the mouse development (Fig. 1B). However, in contrast to the liver, phosphorylation levels of both AMPK and CLIP-170 were significantly elevated at 8 weeks after birth in the heart. Importantly, phosphorylation level of ACC in the heart did not correlate with the increase of AMPK phosphorylation levels after the birth (Fig. 1A). These data suggest that AMPK in the heart might have a distinct role other than metabolism, especially after the birth.

**Figure 1:**
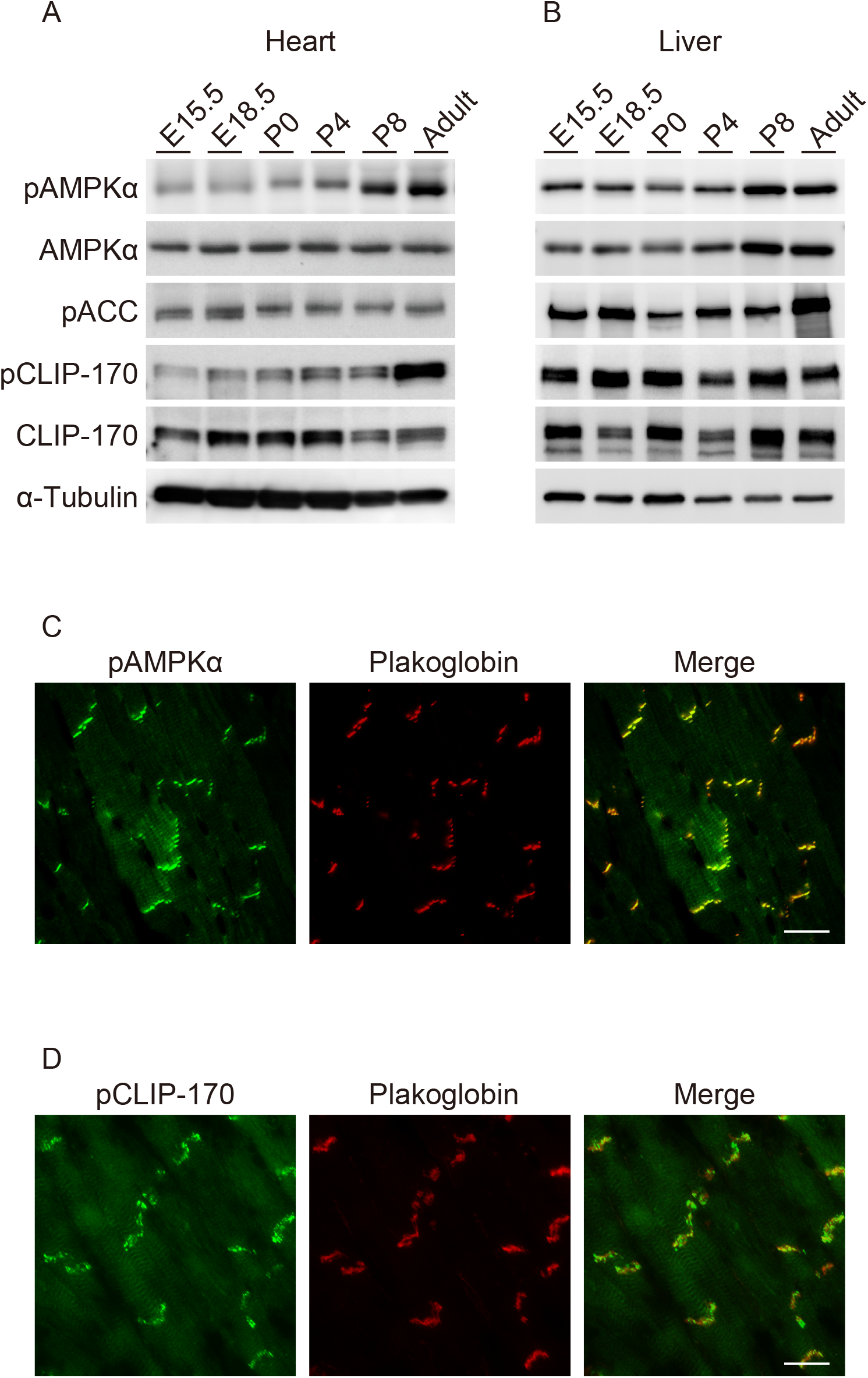
Phosphorylation levels of AMPK significantly increased at the intercalated discs in adult mouse heart together with its substrate CLIP-170. (A, B) Immunoblot analysis of the phosphorylation level of CLIP-170, AMPK and ACC in heart (A) and liver (B) along with the developmental stages. α-Tubulin was used as a loading control. (C, D) Immunostained images of adult mouse heart tissue. These were stained with a phosphorylated AMPK antibody (C, left), a phosphorylated CLIP-170 antibody (D, left) and a plakoglobin antibody (C, D, center). Scale bar, 20 μm (C, D).

Next, we examined the localization of AMPK through immunohistostaining in an adult murine heart. Surprisingly, majority of phosphorylated AMPK was localized at the intercalated discs, which was confirmed through its co-localization with plakoglobin (Fig. 1C). We also confirmed the localization of AMPKβ2 at the intercalated discs using a subunit-specific antibody (Supplemental Fig. 1A). To eliminate non-specific signals in immunohistostaining, we confirmed the localization of AMPK at the cell-cell junctions using the following two methods. In the first method, mCherry-fused AMPKα2, which was expressed in cultured cardiomyocytes, was used to demonstrate AMPK localization at the cell-cell junctions (Supplemental Fig. 1B). Next, we assessed the activity of AMPK at the plasma membrane in cardiomyocytes by using previously described organelle-specific AMPK activity probe, ABKAR [21]. Cardiomyocytes demonstrated significantly higher AMPK activity at the cell-cell junctions compared to HeLa cells (Supplemental Fig. 1C and D). Moreover, the majority of phosphorylated CLIP-170 (Fig. 1D) and LKB1 (Supplemental Fig. 1A), an upstream kinase of AMPK, were found to be localized at the intercalated discs. The intercalated discs are absent in embryonic stages and are eventually formed 7 to 8 weeks after the birth [18]. It is noteworthy that the levels of phosphorylation of AMPK and CLIP-170 assessed by Western blotting were upregulated following the same time course as that of intercalated disc formation (Fig. 1B). Altogether, these findings indicate that AMPK localizes at the intercalated discs in the heart, where it phosphorylates its potential substrate, CLIP-170.

### The localization of AMPK is regulated by contraction of cardiomyocytes

In the heart, proper mechanical stress is essential for maintaining homeostasis, development and cellular function. An individual cardiomyocyte always undergoes mechanical stress due to spontaneous beating, and thus protracted decrease in mechanical stress induces atrophy and cell death in the cardiomyocytes [18]. Cardiomyocytes, or cells in general, possess a mechanical stress sensing system that detects stress or strain. Although the precise molecular mechanism remains unclear, there are several cellular components, including cell membrane and sarcomere-related, which are supposedly involved in mechanical stress sensing [22]. The intercalated disc is one such component, which has been shown to be critical for detecting the mechanical stress generated through myocyte contraction [20].

Therefore, in order to examine whether contraction of cardiomyocytes influences the activity of AMPK at the cell-cell junctions, we performed immunostaining with the anti-phosphorylated AMPKα antibody, which is an indicator of activated AMPK, in rat primary cardiomyocytes. After 2 hours of treatment with MYK-461, a myosin ATPase inhibitor which suppress the contraction of cardiomyocytes, phosphorylated AMPKα signals at the cell-cell junctions were found to be significantly reduced, although the levels of N-cadherin, an adherens junction component, did not change. However, 4 hours after removal of MYK-461, phosphorylated AMPKα reappeared at the cell-cell junctions along with the re-initiation of cardiac beating (Fig 2A). The signal corresponding to a subunit of holoenzyme, AMPKβ2, at the cell-cell junctions depleted within 2 hours of MYK-461 treatment, while the sarcomere-like signal remained unchanged. Similar to the phosphorylated AMPKα, AMPKβ2 reappearance was observed upon removal of MYK-461 (Fig 2B). The localization of connexin43 or plakoglobin, both of which are the components of the cell-cell junctions, was not changed upon MYK-461 treatment (Fig 2C). LKB1 also localized at the cell-cell junctions, but did not disappear upon MYK-461 treatment (Fig 2D). These data indicate that the localization of AMPK in cardiomyocytes is regulated in response to the contraction or mechanical stress of cardiomyocytes.

**Figure 2:**
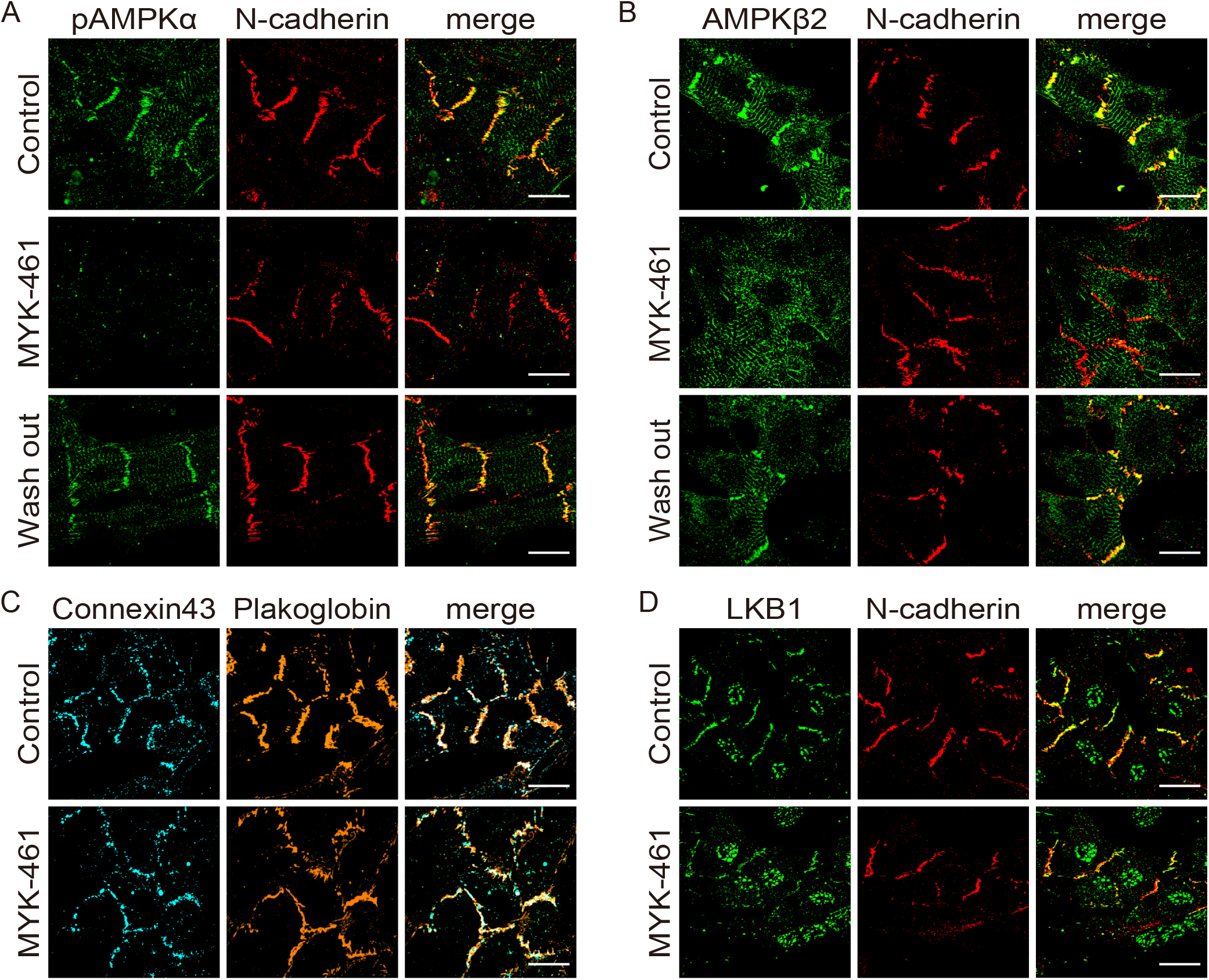
The localization of AMPK was regulated by contraction of the cardiomyocytes. (A, B) Immunostained images of neonatal rat cardiomyocytes 2 hours after treatment with 0.01 % DMSO (Control, upper row) or 4 μM MYK-461(MYK-461, middle row) and 4 hours after washing out of MYK-461 (Wash out, bottom row). These cells were stained with a phosphorylated AMPK antibody, a AMPKβ2 antibody and a N-cadherin antibody. (C, D) Immunostained images of neonatal rat cardiomyocytes 2 hours after treatment with 0.01 % DMSO (Control, upper row) or 4 μM MYK-461(MYK-461, bottom row) and stained with a connexin43 antibody, a plakoglobin antibody, a LKB-1 antibody and an N-cadherin antibody. Scale bar, 20 μm (A-D).

### AMPK regulates MT plus end dynamics through CLIP-170 phosphorylation in cardiomyocytes

As we previously demonstrated that AMPK regulates MT dynamic instability in migrating Vero cells [4], in this study, we assessed the MT dynamics focusing at the intercalated discs in cardiomyocytes. After 3 days of primary culture of cells isolated from rat neonatal heart, cardiomyocytes were found to be connected with the neighboring cardiomyocytes by forming cell-cell junctions composing gap junction proteins, desmosomal proteins and adherens junction proteins, which are the basic constituents of the intercalated disc. However, free plasma membrane without intercellular connections do not possess these junction proteins, suggesting that the heart-specific polarity was established in these primary cultured cells, although they were not completely mature. First, we assessed the intracellular dynamics of CLIP-170 by a time-lapse image analysis in rat neonatal cardiomyocytes. In EGFP-CLIP-170 WT transfected cardiomyocytes, CLIP-170 migrated from the cell’s interior to the periphery with short comet at the plus end of MTs. In cardiomyocytes with rectangular-like shape, the majority of CLIP-170 migrated longitudinally towards the cell-cell junctions (Fig 3A and Movie S1). These results are distinct from the previously published results that CLIP-170 radially moved from the microtubule organizing center to the periphery in migrating cells [4]. After addition of Compound C, an AMPK inhibitor, EGFP-CLIP-170 WT signals became elongated and formed lines, which specifically accumulating at the cell-cell junctions (Fig 3A and Movie S1). To exclude the possibility that the observed change was mediated by non-specific inhibition of other kinases by Compound C, we examined EGFP-CLIP-170 WT dynamics in AMPKα1α2 knockdown (KD) cardiomyocytes using short interfering RNAs (siRNAs). CLIP-170 comets became elongated and accumulated at the cell-cell junctions in AMPK α1α2 KD cardiomyocytes (Fig 3C and D), as observed in Compound C treatment. Moreover, in cardiomyocytes transfected with EGFP-CLIP-170 S311A, which is non-phosphorylatable mutant [4], CLIP-170 also accumulated at the cell-cell junctions with elongated comet as shown in Compound C treated cells, or in AMPK α1α2 KD cells (Fig 3A and B, Supplemental Fig. 2 and Movie S2).

**Figure 3:**
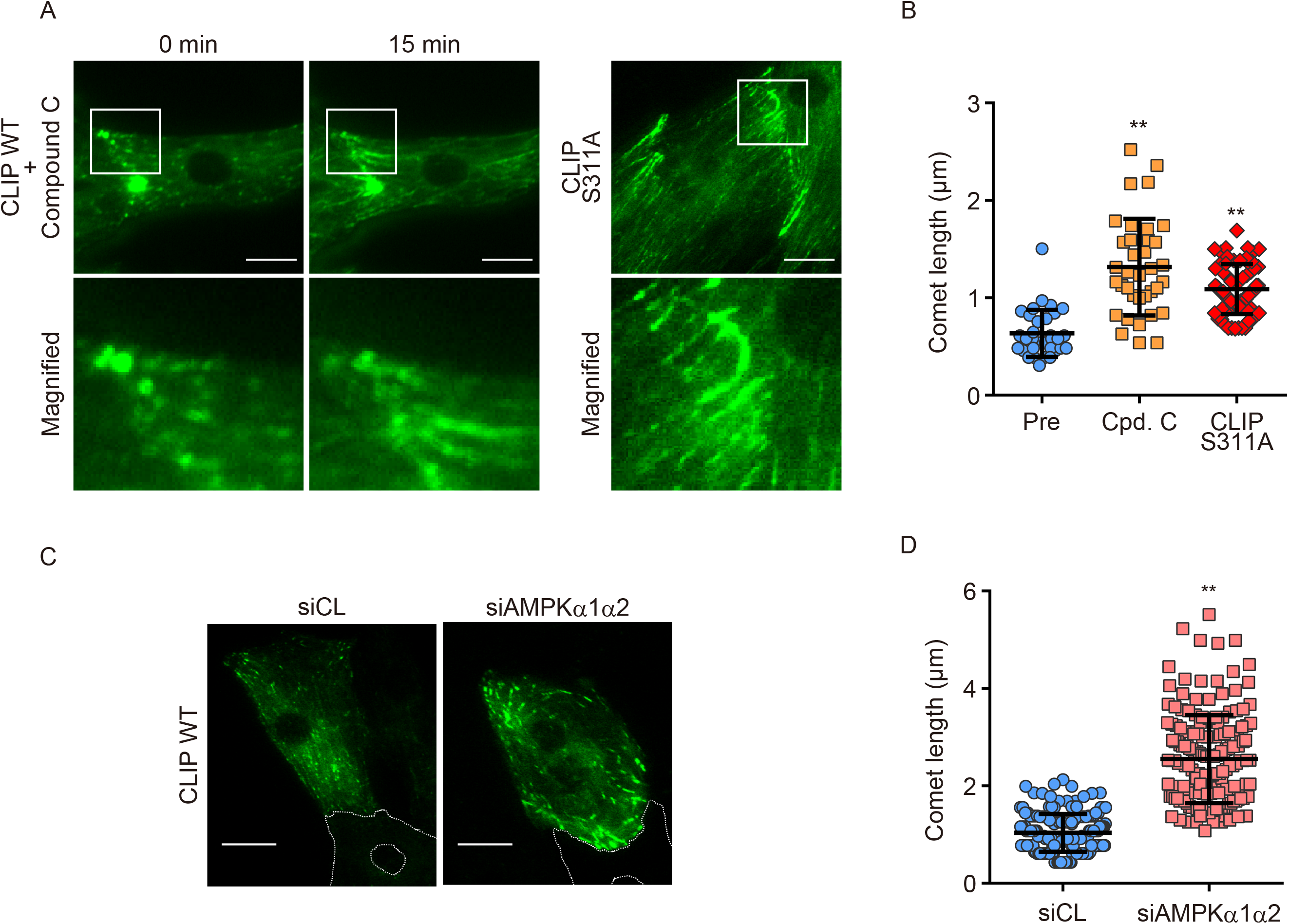
AMPK regulated longitudinal microtubule dynamics through the phosphorylation of CLIP-170 in cardiomyocytes. (A) GFP time-lapse images of neonatal rat cardiomyocytes expressing EGFP-CLIP-170 WT 0 min and 15 min after treatment with 20 μM Compound C (left side panel) and expressing EGFP-CLIP-170 S311A (right side panel). Higher magnification of white square (upper row) showing CLIP-170 migrated longitudinally toward the cell-cell junctions. (B) Scatter plots of a single comet length of neonatal rat cardiomyocytes expressing EGFP-CLIP-170 WT before (Pre) and 15 min after treatment with 20 μM Compound C (Cpd. C) and expressing EGFP-CLIP-170 S311A. Data means ±S.D. Pre: n=32, Cpd. C: n=36, CLIP S311A: n=68, **, P<0.01 versus Pre. (C) GFP time-lapse images of neonatal rat cardiomyocytes expressing EGFP-CLIP-170 WT treated with control siRNA (siCL, left) or siRNA targeting both AMPKα1 and α2 (siAMPKα1α2, right). White dotted lines in the images showed the connected cardiomyocyte not expressing EGFP-CLIP-170 WT. (D) Scatter plots of a single comet length of cardiomyocytes expressing EGFP-CLIP-170 WT treated with control siRNA or siRNA targeting both AMPKα1 and α2. Data means ±S.D. siCL: n=178, siAMPKα1α2: n=180, **, P<0.01 versus siCL. Scale bar, 10 μm (A, C).

These results suggest that MT turnover at the cell-cell junctions is regulated through CLIP-170 Ser 311 phosphorylation by AMPK in cardiomyocytes.

### AMPK controls cell size and shape by regulating MT dynamics through CLIP-170 phosphorylation of in cardiomyocytes

We noticed that a fraction of cardiomyocytes treated with MYK-461 became elongated in the same direction as that of MT migration during the time-lapse image analysis (Fig. 4A). These results directed us to measure the individual cell size of cardiomyocytes treated with MYK-461. Analysis of the imaging data using IN Cell Analyzer revealed that MYK-461 treatment for 2 h significantly increased the cell area of cardiomyocytes (Fig. 4B and C). In order to further elucidate the specific role of CLIP-170 Ser 311 phosphorylation by AMPK, we compared the phenotypes of cardiomyocytes transiently transfected with CLIP-170 WT and two of the Ser 311 mutants of CLIP-170, namely CLIP-170 S311A and CLIP-170 S311D (a phosphomimetic mutant) [4]. The expression of CLIP-170 S311A led to the cell area expansion at baseline, while CLIP-170 S311D mutant had no effect (Fig. 4C). Addition of MYK-461 had no further effect in the cell size in CLIP-170 S311A-expressing cardiomyocytes (Fig. 4C). Conversely, expression of CLIP-170 S311D mutant was refractory to the action of MYK-461 (Fig. 4C). These data suggest that MT dynamics is critical for the maintenance of cell size and shape of the beating cardiomyocytes, which is further dependent on CLIP-170 phosphorylation.

**Figure 4:**
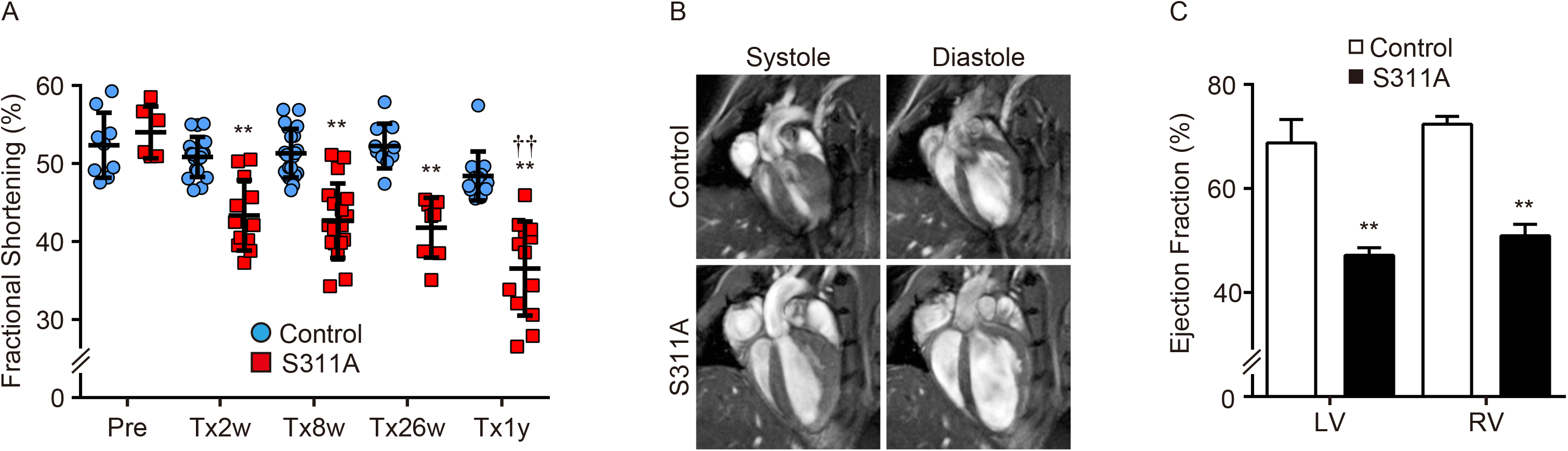
AMPK-CLIP-170 signal at the intercalated disc controlled the cell shape in cardiomyocytes. (A) GFP time-lapse images of neonatal rat cardiomyocytes expressing EGFP-CLIP-170 WT 0 hour, 1 hour and 2 hours after treatment with 4 μM MYK-461. (B) Immunostained images of neonatal rat cardiomyocytes 4 hours after treatment with 0.01 % DMSO (Control, upper row) or 4 μM MYK-461(MYK-461, bottom row). These cells were stained with an α-Tubulin (green) and a plakoglobin (red) antibody. (C) Bar graphs showing the cell size of a cardiomyocyte 2 hours after treatment with or without 4 μM MYK-461. Adenovirus expressing EGFP-CLIP-170 (adCLIP) each mutant was used (WT, S311A, S311D). Data means ±S.D. Control (-/-): n=784, MYK-461: n=939, CLIP WT: n=656, CLIP WT + MYK-461: n=619, CLIP S311D: n=613, CLIP S311D + MYK-461: n=735, CLIP S311A: n=389, CLIP S311A + MYK-461: n=508, **, P<0.01 versus Control, ††, P<0.01 versus CLIP WT, N.S., not significant. Scale bar, 10 μm (A, B).

### Inducible heart-specific *Clip-170* S311A overexpressing transgenic mice exhibit cardiac dysfunction

Thereafter, to investigate the physiological relevance of AMPK-CLIP-170 in vivo, we generated tamoxifen inducible, cardiomyocyte-specific CLIP-170 S311A overexpressing transgenic (TG) mice. In adult CLIP-170 S311A flox/+; MerCreMer+/− mice, we successfully confirmed CLIP-170 S311A protein expression in cardiac muscle from these mice 2 weeks after the initiation of tamoxifen administration. We established two lines of CLIP-170 S311A TG mice (Supplemental Fig. 3), we present the data of line 3 hereafter. We found that they were phenotypically similar. To check the influence of overexpression of CLIP-170 S311A in cardiac function, we performed serial echocardiography measurements. Two weeks after tamoxifen induction, CLIP-170S311A TG mice showed a mild, but significant decline in fractional shortening, known as an indicator of cardiac function, compared to the control mice. Over 1 year after tamoxifen administration, CLIP-170 S311A TG mice exacerbated cardiac dysfunction (Fig. 5A). To further analyze both ventricles, we performed cardiac MRI in these mice. Both left and right ventricle were found to be significantly dilated and with reduced contraction significantly in CLIP-170 S311A TG mice (Fig. 5B and C). The histological assessment of CLIP-170 S311A TG mice after 3 months of tamoxifen treatment showed that there was no tissue degeneration, however the cardiac function was impaired. However, CLIP-170 S311A TG mice over 1 year after tamoxifen treatment showed significant tissue fibrosis compared to the control (Fig. 6A and B). Immunohistostaining revealed enhanced accumulation of tubulin in CLIP-170 S311A TG mice (Fig. 6C). Next, we checked the individual cell size in the heart of CLIP-170 S311A TG mice by wheat germ agglutinin (WGA) staining, a lectin that stains cell membrane. CLIP-170 S311A TG mice showed that the length of individual cells was significantly elongated in a long axis direction compared to the control (Fig. 6D and E).

**Figure 5:**
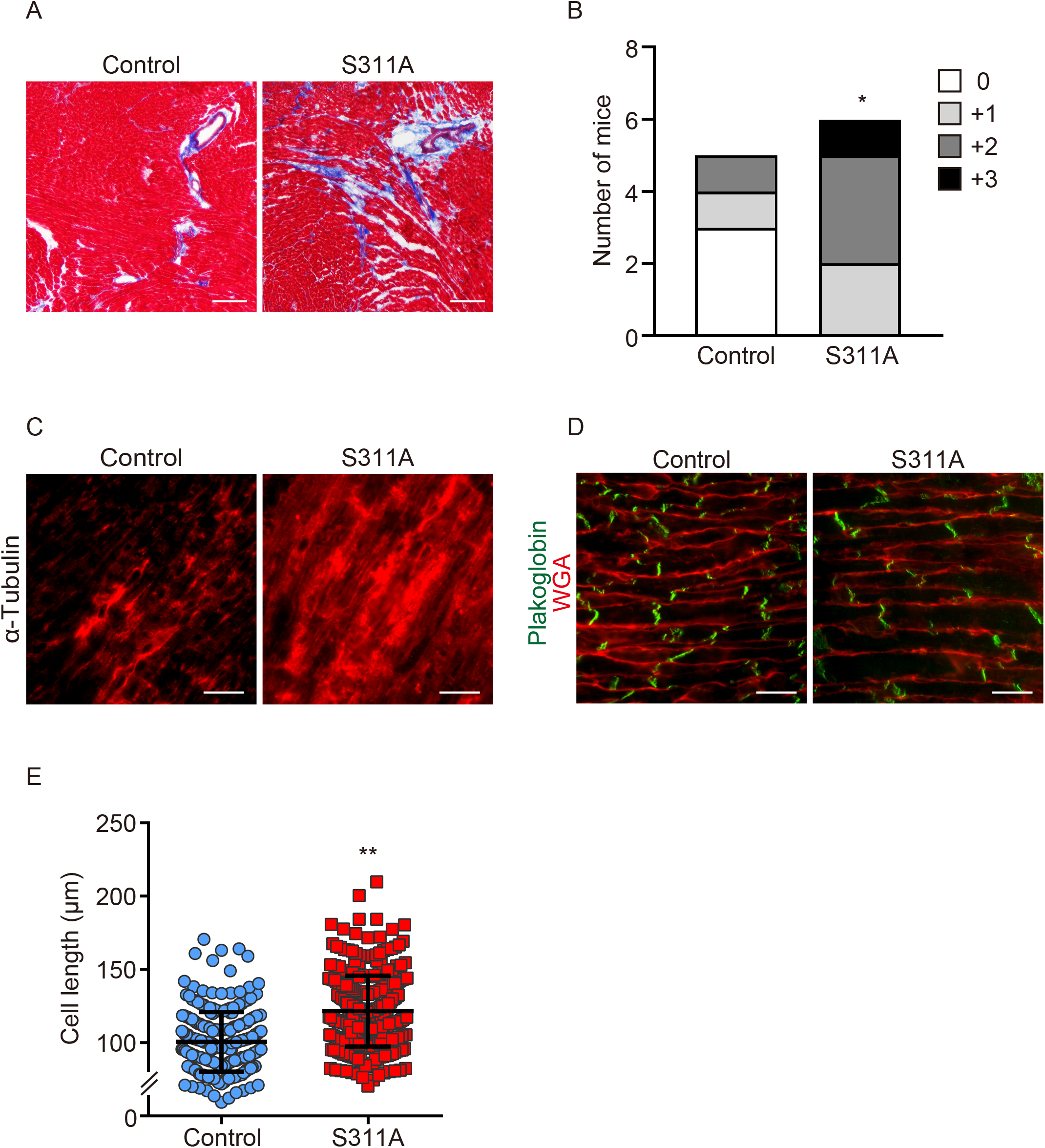
Inducible heart-specific CLIP-170 S311A overexpressing transgenic mouse shows cardiac dysfunction. (A) Scatter plots of echocardiographic parameter (fractional shortening) of individual CLIP-170 S311A overexpressing mice and control mice before (Pre), 2 weeks (Tx2w), 8 weeks (Tx8w), 26 weeks (Tx26w) and over 1 year (Tx1y) after the tamoxifen induction. Data means ±S.D. Pre Control: n=9, Pre S311A: n=6, Tx2w Control: n=18, Tx2w S311A: n=12, Tx8w Control: n=21, Tx8w S311A: n=18, Tx26w Control: n=10, Tx26w S311A: n=9, Tx1y Control: n=12, Tx1y S311A: n=13, **, P<0.01 versus each Control, ††, P<0.01 versus Pre S311A. (B) Representative long axis four-chamber cardiac magnetic resonance images of CLIP-170 S311A overexpressing mice and control mice over 1 year after the tamoxifen induction. Left column showed systole images and right column were Diastole images. (C) Bar graphs showing the ejection fraction of CLIP-170 S311A overexpressing mice and control mice over 1 year after the tamoxifen induction. LV, left ventricular, RV, right ventricular. Data means ±S.D. Control: n=3, S311A: n=3, **, P<0.01 versus each Control.

These data indicate that the physiological CLIP-170 phosphorylation by AMPK at the cell-cell junctions is important for homeostatic MT dynamics, thereby maintaining the cell shape and cardiac function of cardiomyocytes.

## Discussion

In this study, we demonstrated that cardiac MTs are dynamic and undergo constant turnover at the intercellular junctions, as revealed by time-lapse image analysis. Most of the AMPK in beating cardiomyocytes is localized at the cell-cell junctions where it phosphorylates CLIP-170, thereby regulating MT turnover. It is noteworthy that the myosin ATPase inhibitor, MYK-461, prevents the localization of AMPK at the cell-cell junctions, and this effect is reversed by the removal of MYK-461. The inhibition of CLIP-170 phosphorylation or MYK-461 treatment, which alters subcellular localization of AMPK, results in an increase in the cell area of cardiomyocytes. Moreover, the heart-specific CLIP-170 S311A overexpressing TG mice showed the elongation of cardiomyocytes with accumulated MTs, which ultimately resulted in a progressive decline in cardiac contractile function.

Cardiac MTs have drawn attention in cardiovascular research as a modulator of intracellular stiffness. Accumulation of MTs has been associated with human heart failure and rodent models of hypertrophy, myocardial ischemia-reperfusion, catecholamine-induced myocardial injury, heart failure [11,12,15,16,23]. Recently, Prosser lab elegantly revealed that tubulins with increased post-translational modification (PTM) confer mechanical resistance to contraction and regulate the viscoelastic properties of myocyte [15]. Detyrosinated MTs are found to be elevated in human or rodent heart failure models, suggesting that PTM of cardiac MTs has a relevance in the pathology of heart diseases, however, the mechanisms that are responsible for the accumulation of tubulins or PTM in heart diseases are not well understood.

Cardiac MTs have been recognized as static constituents of the cellular cytoskeleton, especially in highly differentiated cells, including cardiomyocytes. Our findings suggest that MTs in cardiomyocytes are rather dynamic and their turnover is regulated by CLIP-170 phosphorylation, which is mediated by specially localized AMPK in the intercalated discs. CLIP-170 is an MT plus end tracking protein (+TIPs), which in involved in maintaining the equilibrium of MT dynamics towards extension rather than catastrophe. Once CLIP-170 is phosphorylated by the upstream kinases, for instance, AMPK in this case, its affinity to MT is decreased [8]. Therefore, it is reasonable that the affinity of CLIP-170 to MTs is decreased upon phosphorylation at the intercalated discs, where MT polymerization is supposed to terminate. We previously demonstrated that the inhibition of CLIP-170 phosphorylation by AMPK increases detyrosinated MTs, leading to accumulation of MTs [4]. In this study, CLIP-170 S311A TG mice showed decreased cardiac function upon accumulation of MTs, which is often found in heart failure models. It is thus possible that the perturbation of CLIP-170 phosphorylation or inhibition of AMPK at the intercalated discs in cardiomyocytes is involved in the pathogenesis of heart diseases, which could be mediated by an increase in detyrosinated MTs [16]. It has been shown that AMPK is activated in a rodent and human heart failure samples [24,25], although another study showed decreased activity in a rat spontaneous hypertensive model [26]. Considering the fact that we revealed specific subcellular localization of AMPK at the intercalated discs in an adult murine heart, it will be interesting to assess AMPK activity in each subcellular compartment using heart failure models in future studies.

Inhibition of cardiac contraction by MYK-461 or inhibition of phosphorylation, as shown by using the CLIP-170 S311A mutant, surprisingly led to increase in cell size. It is noteworthy that cardiomyocytes in the CLIP-170 S311A TG mice became elongated and showed an increase in the aspect ratio of individual cells. A previous report demonstrated that cardiomyocytes adapt to the elasticity of extracellular matrix and modulate their cell shape and length in order to maximize their systolic performance, and also that MT is a key component of cardiomyocytes [17]. In other words, cardiomyocytes have an intrinsic mechanism to adjust their cell shape or aspect ratio through change in MT polymerization. Therefore, it is likely that perturbation in MT dynamics leads to failure in maintaining the optimal aspect ratio of the cardiomyocytes, resulting in decreased contraction, which might be the consequence of what we observed in the CLIP-170 S311A TG mice. Elongation of cardiomyocytes is often observed in heart failure models or human end-stage heart failure [17,27–29]. And it is considered a part of the vicious cycles in the pathogenesis of heart failure. Thus, maladaptation of cardiomyocyte cell shape through altered MT dynamics could be a domain that needs further consideration in future cardiovascular research. From another view point, there are many reports suggesting that MTs play an important role as an endogenous factor regulating the contractile force of cardiomyocytes through modulating intracellular stiffness [15,16]. Therefore, in our CLIP-170 S311A TG mice, the accumulation of MTs themselves may explain the reduced contractility observed in these mice.

Our results suggest that the activity of the AMPK, which is specifically localized at the intercalated discs, does not correlate with energy metabolism (sensing AMP/ATP ratio), but it does correlate with mechanical stress. Although the localization of AMPK in other tissues have not been fully examined yet, it was reported that AMPK activity is involved in cell-cell junctions in lung epithelium and alveolar development in response to repeated respiration-induced physical stretching [30]. In fact, AMPK and its upstream LKB1 ortholog have been found to exist in animals lower than mammalian order, however, there is no such report that these enzymes are involved in energy metabolism [9]. In budding yeast, the SNF1 complex corresponding to the AMPK ortholog has been shown to be activated upon glucose-starved state, but is not allosterically activated by AMP [31]. The AMP/ADP-sensing property of AMPK is considered to have been acquired, at least in mammals [32], suggesting that the ancestral regulation and/or function of AMPK might be different. Certainly, it has been reported that fructose-1,6-diphosphate and aldolase mediate glucose sensing by AMPK localized in lysosomes occurs without being activated by AMP/ADP [33]. It is possible that the mechanosensing property of AMPK is an evolutionary descendant of the ancestral AMPK, and it may be found in other tissues in mammals or other organisms.

In this study, we demonstrated that AMPK localization is dynamically regulated by the beating of cardiomyocytes, suggesting the involvement of mechanical stress. LKB1 is one of the upstream kinases of AMPK, which is constitutive active. Interestingly, we found that LKB1 was co-localized at the cell-cell junctions along with AMPK; however, MYK-461 treatment did not influence its localization (Fig. 2D). Therefore, it seems subcellular localization of AMPK is crucial for its mechanical signaling sensing. AMPK is a multifunctional kinase with multiple substrates and cellular outcomes [21]. Thus, it is conceivable that specific subcellular localization enables AMPK to play distinct roles in different cell types. AMPK has been shown to be activated by mechanical stretch in skeletal muscle cells [34], or in the lung epithelial cells [35]. Additionally, LKB1 has been shown to be recruited to the cadherin adhesion complex in response to force and thereby activates AMPK in epithelial cells [36,37]. From these reports and our findings, it is possible that AMPK is involved in mechanotransduction; however, LKB1-AMPK regulation in response to mechanical stress could depend on types of cells or stimulation. We were not able to reveal the molecular mechanism involved in the translocation of AMPK in response to mechanical signaling in cardiomyocytes, and thus further investigation is required to be performed in future studies.

The number of patients with chronic heart failure has increased globally, and there is an urgent need to develop effective treatment strategies for heart failure with novel mechanisms of action. Previous studies have provided strong evidences that cardiac MTs play crucial role in the pathogenesis of heart failure [16]. Thus, it is extremely important to understand the molecular mechanism that regulates MT dynamics in the physiological and pathological heart conditions so that the molecular targets can be identified to establish effective treatment strategies.

## Methods

### Plasmids and viral constructs

To create adenoviral vectors expressing full-length mouse CLIP-170 WT/S311A/S311D fused with enhanced green fluorescent protein (EGFP) [4,38], the corresponding cDNAs were subcloned into pENTR for further Gateway recombination in adenoviral expression plasmids, pAdCMV/V5/DEST (Invitrogen). Recombinant adenoviral vectors were produced and purified using HEK293A cells according to manufacturer’s protocol (ViraPower Adenoviral Expression System; Invitrogen, AdenoPACK 20, Vivapure; Sartorius AG).

Generation of rabbit polyclonal antibodies specific for the phosphorylated S311 of CLIP-170. A phospho-specific polyclonal antibody to CLIP-170 (Ser 311) was generated by Scrum Inc. as follows. Ser-phosphorylated or non-phosphorylated peptides surrounding S311 (amino acids 305–316, SLKRSP(pS)ASSLS) was synthesized. Rabbits were immunized 5 times with the keyhole limpet hemocyanin–phosphopeptide conjugates mixed with Freund’s complete adjuvant, and bled 7 days after the last immunization. Phosphopeptide-reactive antibody was captured and eluted by a column containing phosphopeptide-conjugated sepharose. Then, non-specific fraction was removed using a column containing non-phosphorylated peptides. Specific reactivity with the targeted phosphoserine sequence was confirmed by an ELISA in which phosphorylated or non-phosphorylated peptides were coated.

### Cell culture, plasmid transfection, and siRNA

Primary cultures of neonatal cardiomyocytes were prepared from 1- to 3-day-old Wistar rats as described previously [39,40]. Briefly, harvested hearts were incubated in 0.25% trypsin/EDTA (Sigma) at 4°C overnight and then digested with collagenase type II (Worthington). The cardiomyocyte fraction was collected after differential plating for 70 min at 37°C, counted, and seeded onto plates or collagen-coated glass-bottom dishes. Cardiomyocytes were cultured in DMEM (Sigma-Aldrich) supplemented with 10% FBS (Sigma-Aldrich), penicillin and streptomycin (Gibco) at 37°C in a 5% CO_2_ atmosphere at constant humidity.

For the time-lapse imaging, cardiomyocytes seeded on collagen-coated 35-mm glass dishes were transfected with adenovirus expressing EGFP-CLIP-170 WT/S311A/S311D at 48 hours after isolation and observed at 24 hours after transfection.

To knockdown AMPKα1 and α2, cardiomyocytes were transfected with siRNAs (Silencer® Select siRNA; AMPKα1 siRNA ID: s134808 (30 nM), AMPKα2 siRNA ID: s134962 (10 nM), Thermo Fisher Scientific) using lipofectamine RNAi MAX (Invitrogen) at 3 hours after isolation.

As a negative control, cells were transfected with siControl Non-Targeting siRNA#1 (B-bridge).

Isolation of mRNA and protein experiments were performed at 72 hours after transfection. For immunostaining, the same procedures of siRNA transfection were performed in one-fifth scale on Lab-Tek Chamber Slides (nunc).

### Immunoblotting

Protein concentration was determined using the BCA protein assay kit (Thermo Fisher Scientific). Equal amounts was fractionated by SDS-PAGE, transferred to a PVDF membrane by electroblotting, and processed for Immunoblotting, as described elsewhere [4]. Blots were probed with the appropriate specific antibodies (Anti-AMPKα, 1:1000; Anti-pAMPK, 1:1000; Anti-pACC, 1:1000; Anti-CLIP170, 1:1000; Anti-pCLIP170, 1:1000;

Anti-α-Tublin, 1:5000 dilution), followed by secondary antibodies, and developed by ECL chemiluminescence (GE Healthcare).

### Immunostaining and immunohistology

Cardiomyocytes were seeded on a collagen-coated 35-mm glass dishes (Iwaki). After cells firmly attached to the dish, they were washed once with warm PBS and fixed with methanol for 15 min at 20°C. Next, the cells were permeabilized with 0.1% Triton X-100 in PBS for 5 min at room temperature and then blocked with 1% BSA and 5% goat serum for 15 min at room temperature. Samples were immunostained with primary antibodies (1:200 in 1% BSA and 5% goat serum, overnight). The next day, for secondary reactions, species-matched Alexa Fluor 488. or Alexa Fluor 568. or Alexa Fluor 647-labelled secondary antibody was used (1:400 in 1% BSA, 30 min). Fluorescence images of EGFP, Alexa Fluor 488, Alexa Fluor 546, Alexa Fluor 568 and Alexa Fluor 647 were recorded using an Olympus FV1000-D confocal laser scanning microscope (Olympus Corporation) equipped with a cooled charge-coupled device CoolSNAP-HQ camera (Roper Scientific, Tucson, AZ, USA) and a PLAPO ×60 oil-immersion objective lens.

The mouse heart was perfused with ice-cold PBS, then removed, cut and embedded in O.C.T. compound and frozen in isopentane chilled in liquid nitrogen. The frozen tissue sections (7-10 μm thick) were fixed with MeOH at −20°C for 10 min. After permeabilization with PBS containing 0.1% of TritonX100 for 5 minutes at the room temperature, and the non-specific antibody-binding sites were pre-blocked with the blocking buffer (PBS plus 5% of goat serum and 1% of bovine serum albumin). The primary antibodies were applied overnight at 4°C. After rinsing 3 times for 5 minutes in PBS, the sections were next incubated with appropriate fluorophore-conjugated secondary antibodies and 4’,6-diamidino-2-phenylindole (DAPI) in the blocking buffer for 30 min at room temperature. The primary antibodies used in this study are as follows;

| Primary antibodies | Host | dilution |  |
| --- | --- | --- | --- |
| phosphorylated AMPKα | rabbit | 1/100 | Cell signaling |
| Connexin43 | rabbit | 1/200 | abcam |
| Plakoglobin | guinea pig | 1/400 | PROGEN |
| AMPKβ2 | rabbit | 1/100 | HPA |
| N-cadherin | mouse | 1/200 | Santa Cruz |
| α-actinin | mouse | 1/200 | abcam |
| AMPKα | rabbit | 1/500 | Cell Signaling |
| CLIP-170 | mouse | 1/500 | HPA |
| α-Tubulin | rabbit | 1/1000 | Cell Signaling |
| phosphorylated CLIP | rabbit | 1/500 | Custom made |
| phosphorylated ACC | rabbit | 1/1000 | Cell Signaling |
| LKB1 | rabbit | 1/200 | Cell signaling |

### Time-lapse imaging and tracking

The time-lapse imaging was performed as described previously [4]. Briefly, fluorescence images were recorded by using the same experimental equipment and software described in the section above. An EGFP image was obtained every second through a U-MNIBA2 filter (Olympus), which had a 470–495 excitation filter and a 510–550 emission filter. To achieve high temporal resolution, we had to limit the exposure time to 200 msec. CLIP-170 kinetics were analyzed on 16-bit depth images after subtraction of the external background. We measured the fluorescence intensity values within the line of 1 pixel in width along the CLIP-170–EGFP tracks over time. We determined the beginning of a comet as the point at which fluorescence intensity showed a rapid rise and the end of a comet as the point at which fluorescence intensity reached baseline. MetaMorph was used to convert a series of time-lapse images to video format and obtain tracking images in this analysis.

### The quantitative evaluation of surface area of cardiomyocytes

We performed the quantitative evaluation of surface area of cardiomyocytes by using IN Cell Analyzer (GE healthcare). αActinin positive cardiomyocytes with nuclei (Hoechst) were selected and cell bodies were decided on the basis of intensity. The mean cell surface area of each cardiomyocyte was calculated as a numerical value.

### Animals

All animal experiments were approved by the Animal Research Committee of Osaka University, and were performed in accordance with institutional guidelines.

### Generation of cardiomyocytes specifically CLIP-170 S311A-overexpressing mice

CLIP-170 S311A was subcloned into CAG-loxp-CAT-loxp vector, and transgenic strain by pronuclear injection in mouse zygotes. Mice bearing the CLIP-170 S311A flox/+ allele were crossed with a transgenic line expressing Cre recombinase under the control of the α-myosin heavy chain promoter (MerCreMer: provided by Dr Molkentin) in a tamoxifen-inducible cardiomyocyte-specific manner to produce CLIP-170 S311A flox/+; MerCreMer+/− mice [41]. CLIP-170 S311A flox/+; MerCreMer−/− littermates were used as age-matched controls. The CLIP-170 flox/+; MerCreMer+/− mice were genotyped by PCR using primers for CAT gene and Cre recombinase. In adult CLIP-170 S311A flox/+; MerCreMer+/− mice treated with tamoxifen for 6 days (daily dose of 20 mg kg^−1^), Cre recombination was confirmed by checking CLIP-170 S311A protein levels in cardiac muscle from these mice 14 days after the onset of tamoxifen treatment. We established and analyzed 2 lines of S311A mice (Supplementary figure 3). We presented the data of line 3; their phenotypes were basically similar.

### Cardiac MRI

Serial MRI was conducted using a horizontal 7.0 T Bruker scanner (BioSpec 70/30 USR, Bruker Biospin). All MRI experiments were performed under general anaesthesia using 1%– 2% isoflurane administered via a mask covering the nose and mouth of the animals. Respiratory signals, body temperature, and heart rate were monitored using a physiological monitoring system (SA Instruments, Inc.). Body temperatures were continuously maintained at 36.0 + 0.5°C by circulating water through heating pads throughout all experiments [42]. The center of the imaging slice was carefully positioned at the mouse hearts. First, a three-plane sequence was performed for the definition of slice orientation using self-gated cine imaging with navigator echo. Next, six consecutive scans of the short axis from the apex to the base of hearts were obtained in the long axis four-chamber and long axis two-chamber views. These eight scans were used for fast low-angle shots with navigator echo (IntraGate, Bruker) using the following parameters: repetition time/echo time = 6.0/2.2 ms, flip angle = 10 degrees, field of view = 2.56 × 2.56 cm, matrix = 256 × 256, slice thickness = 1.0 mm, number of repetitions = 300, four concomitant slices covering the whole heart from the apex to base, 10 phases per cardiac cycle, expected heart rate = 400 beats per minute (bpm), expected respiratory rate = 60 bpm, in-plane resolution per pixel = 100 μm, acquisition time = 3 minutes 50 seconds per scan, total acquisition time = approximately 35 min, and a total anesthesia time = approximately 40 min.

### MRI data analysis

In short-axis images, end-diastolic and end-systolic frames were selected according to maximal and minimal ventricular diameter. The epicardial border was manually outlined and the LV cavity was segmented in both frames using software ImageJ. The respective volumes were calculated as the area of each compartment multiplied by the slice thickness (1.0 mm). Based on end-systolic and end-diastolic volumes [ESV (μl) and EDV (μl), respectively), all parameters characterizing cardiac function, such as stroke volume [SV (μl) = EDV - ESV] and ejection fraction [EF (%) = SV/EDV] were calculated.

### Statistical analyses

Data are expressed as means ± S.D. The two-tailed Student’s t-test was used to analyze differences between two groups. Differences among multiple groups were compared by one-way ANOVA, followed by a post hoc comparison using the Tukey method with Prism 6 (GraphPad). For the histological assessment of CLIP-170 S311A mutant mice, comparison was made by Cochran-Armitage trend test. P <0.05 was considered statistically significant.

## Acknowledgments

We also thank Ms Gion, Shingu, Takata for their technical assistance. This study was supported by grants from the Ministry of Education, Culture, Sports, Science and Technology of Japan (15K08271, 18K08105), Mochida Memorial Foundation for medical and pharmaceutical research, Suzuken Memorial Foundation.

## Author contributions

YS conceived and designed the research; S. Yashirogi conducted most of the experiments in this study; TK, TN, YN, TSM, SS, H. Kioka, YM, IY, KY contributed data collection, analysis and figure preparation. HU, OY provided the resources and experimental advice. YN, OT, S. Yamazaki, H. Kato, KY, ST, discussed the data from expert knowledge; S. Yashirogi, S. Yamazaki and YS wrote the manuscript.

---

**Figure 6: CLIP-170 S311A overexpressing transgenic mice showed elongation of the cardiomyocytes with MT accumulation.** (A) Masson’s trichrome staining of the heart of CLIP-170 S311A overexpressing mice and control mice over 1 year after the tamoxifen treatment. (B) Semi-quantitative scaling of cardiac fibrosis in CLIP-170 S311A overexpressing mice and control mice over 1 year after the tamoxifen treatment. Scaling class, 0, only perivascular fibrosis (white), +1, little interstitial fibrosis (light grey), +2, local interstitial fibrosis more than +1 (grey), +3, extensive interstitial fibrosis (black). *, P<0.05 versus Control. (C) Immunostained images with an α-Tubulin antibody of CLIP-170 S311A overexpressing mice heart and control mice heart over 1 year after the tamoxifen treatment. (D) Representative immunostained images of *Clip-170* S311A overexpressing mice heart and control mice heart over 1 year after the tamoxifen treatment. These were stained with WGA (red) and a plakoglobin antibody (green). (E) Scatter plots of a single cell length of CLIP-170 S311A overexpressing mice heart and control mice heart over 1 year after the tamoxifen treatment. Data means ±S.D. Control, n=259, S311A, n=326, **, P<0.01 versus Control. Scale bar, 100 μm (A), 20 μm (C, D).

